# The Arabidopsis Target of Rapamycin (TOR) kinase regulates ammonium assimilation and glutamine metabolism

**DOI:** 10.1101/2022.12.09.519783

**Authors:** Camille Ingargiola, Isabelle Jéhanno, Céline Forzani, Anne Marmagne, Justine Broutin, Gilles Clément, Anne-Sophie Leprince, Christian Meyer

## Abstract

In Eukaryotes, Target of Rapamycin (TOR) is a well conserved kinase that controls cell metabolism and growth in response to nutrients and environmental factors. Nitrogen (N) is an essential element for plants and TOR functions as a crucial N and amino acid sensor in animals and yeast. However, the knowledge on the connections between TOR and the overall N metabolism and assimilation in plants is still limited. In this study, we investigate the regulation of TOR in Arabidopsis by the N source as well as the impact of TOR deficiency on N metabolism. Inhibition of TOR globally decreases ammonium uptake while triggering a massive accumulation of amino acids such as Gln, but also of polyamines. Coherently, TOR complex mutants were found to be hypersensitive to Gln. We also show that the glutamine synthetase inhibitor glufosinate abolishes Gln accumulation resulting from TOR inhibition and improves the growth of TOR complex mutants. These results suggest that a high level of Gln contributes to the reduction in plant growth resulting from TOR inhibition. Glutamine synthetase activity was reduced by TOR inhibition while the enzyme amount increased. In conclusion our findings show that the TOR pathway is intimately connected to N metabolism and that a decrease in TOR activity results in a glutamine synthetase-dependent Gln and amino acids accumulation.

**One sentence summary:** The conserved Target of Rapamycin (TOR) kinase is an important sensor and regulator of the nitrogen metabolism and here we show that inhibiting this kinase affects ammonium uptake and results in Gln accumulation in a glutamine synthetase-dependent manner.

## INTRODUCTION

Sensing available nutrients and efficiently assimilating them is a challenge common to all living organisms for optimal growth and adaptation to changing environments. The tight control of cellular processes involved in nutrients assimilation by different levels of regulatory steps is crucial for maintaining metabolic homeostasis within the cells and between organs (Efeyan et al., 2015). Indeed, when nutrients are abundant, high levels of protein synthesis support cell division and differentiation. By contrast, growth has to stopped in nutrient poor conditions to preserve energy while nutrient stores need to be remobilized to sustain basic metabolism. Nitrogen (N) is an essential macro-element for plants and its availability is important for growth and thus for crop yield (O’Brien et al., 2016; Fichtner et al., 2021). Plant roots can take up N from the soil and plants mainly use inorganic N sources such as nitrate (NO_3_) or ammonium (NH_4_) (Bloom et al., 1992, for review see Kiba and Krapp, 2016). Nitrate can be stored in the vacuole, assimilated directly in the roots or transported to the shoots for its assimilation into nitrite, ammonium and finally glutamine (Krapp et al., 2015; Vidal et al., 2020). Several enzymes are involved in this assimilation: Nitrate and Nitrite Reductases (NR and NiR), Glutamine Synthetase (GS) which converts ammonium and glutamate into glutamine, and Glutamine Oxoglutarate Aminotransferase (GOGAT) which produces two molecules of glutamate by trans-amination of alpha-ketoglutarate. Consequently, GS and GOGAT enzymes are also involved in ammonium detoxification mechanisms (Guan et al., 2016). Assimilation of N is also needed for the synthesis of other N-containing molecules, like other amino acids, polyamines, nucleotides, secondary metabolites or hormones.

In Eukaryotes, the TOR (Target Of Rapamycin) protein kinase is a conserved and important regulatory element controlling many aspects of C and N metabolism in relation with central cellular processes like mRNA translation, cell proliferation, and autophagy (Wullschleger et al., 2006; Gonzalez and Hall, 2017; Brunkard, 2020; Ingargiola et al., 2020; Liu and Sabatini, 2020). Belonging to the Phosphatidylinositol Kinase-related Kinase (PIKK) family, this large Ser/Thr kinase is involved in two protein complexes in Yeast and Mammals. In the TORC1 complex, TOR interacts with LST8 (Lethal with Sec Thirteen 8) and KOG1 (Kontroller of Growth protein 1)/RAPTOR (Regulatory-Associated Protein of TOR) (Adami et al., 2007) whereas in the TOR complex 2 (TORC2), TOR interacts with LST8 and AVO3 (Adheres Voraciously to TOR2 protein3)/RICTOR (Rapamycin-Insensitive Companion of mTOR).

All photosynthetic organisms contain genes coding for TOR, LST8 and RAPTOR proteins but there is so far no evidence for the presence of a TORC2 complex (Maegawa et al., 2015; Dobrenel et al., 2016a; Perez-Perez et al., 2017). LST8 binds the TOR kinase domain to stabilize the complex (Moreau et al., 2012; Aylett et al., 2016) and RAPTOR interacts with the HEAT repeats of the Arabidopsis TOR kinase (Mahfouz et al., 2006). Both proteins probably regulate the docking and phosphorylation of specific TOR substrates. While *tor* mutations are embryo lethal at an early stage of plant development, *lst8* and *raptor* mutants are viable, but present many developmental and metabolic alterations (Menand et al., 2002; Anderson et al., 2005; Deprost et al., 2005; Moreau et al., 2012; Salem et al., 2018). For example, *lst8* mutants contain higher concentrations of almost all free amino acids but this is particularly striking for glutamine (Gln) which accumulates to high amounts (Moreau et al., 2012; Forzani et al., 2019). Recently, inactivation of the dual specificity Tyr/Ser YAK1 (Yet Another Kinase 1) kinase was identified as a strong suppressor of TOR deficiency in Arabidopsis (Barrada et al., 2019; Forzani et al., 2019). The YAK1 kinase interacts with RAPTOR and is a direct TOR substrate (Forzani et al., 2019). Interestingly, inhibition of the YAK1 kinase in the *lst8* background restores growth and strongly decreases glutamine accumulation (Forzani et al., 2019). YAK1 was originally found in yeast where it phosphorylates and regulates transcription factors like Hsf1 and Msn2/4 involved in the responses to stress and nutrient starvation (Lee et al., 2008).

Like for *lst8* mutants, mutations in RAPTOR (Salem et al., 2018) or TOR inhibition by silencing in Arabidopsis (Ren et al., 2012; Caldana et al., 2013) also trigger an increase in amino acids contents. The same trend has also been observed after rapamycin (TOR inhibitor) treatment of *Chlamydomonas reinhardtii* (Mubeen et al., 2018). This amino acid accumulation could be due to an increased recycling of N-containing molecules by autophagy, a process that is activated by TOR inhibition and/or to a decrease in mRNA translation (Liu and Bassham, 2010; Diaz-Troya et al., 2011; Soto-Burgos and Bassham, 2017). In Chlamydomonas the build-up of amino acids has been ascribed to an increased assimilation of exogenous N since this accumulation was reduced when cells were beforehand starved for C or N nutrients but not when treated with inhibitors of translation or proteolysis (Mubeen et al., 2018).

In yeast, TOR also has a strong impact on Gln metabolism by repressing the nuclear localization of the central Gln3 transcription factor which is the regulator of ammonium assimilation (Crespo et al., 2002). Consistently, growth on poor or limited N sources inhibits TOR which results in Gln3 activation (Beck and Hall, 1999; Tate et al., 2021). In animals Gln was shown to be a strong inducer of TOR activity through glutamine catabolism (Yuan et al., 2015; Villar et al., 2015; Tanigawa and Maeda, 2017). Indeed, cancerous cells often have an increased glutamine catabolism, which promotes high TOR activity, to fuel their high needs for energy and nutrients (Martinez-Outschoorn et al., 2017).

More generally, several studies demonstrated in yeast and mammals that TOR activity is regulated by nutrient availability and amino acids. As an example, Stracka et al. showed by following Sch9 phosphorylation (a direct TOR target), that N source acts on TOR activity in yeast. The authors determined two N source categories, one “high-end” (like glutamine, ammonium, asparagine) and one “low-end” (serine, branched amino acids and hydrophobic amino acids), the first category inducing TOR activity while poor N sources inhibit TOR (Stracka et al., 2014). In the same way, Meng et al., demonstrated that in human embryonic kidney HEK293A cells, amino acids activate mTORC1 and promote its localization to the lysosome (Meng et al., 2020). Indeed, in animals and yeast, TORC1 is stimulated following recruitment to the lysosomal/vacuolar surface by amino acid activated V-ATPase through the RagGTPase/RagulaTOR complex (Liu and Sabatini, 2020). As in yeast and mammals, TOR activity in plants is also regulated by nutrient availability, especially sugar, sulfur and amino acids (for review Ingargiola et al., 2020). In stress conditions or during C starvation, SnRK1 (Snf1-Related Kinase 1) is activated, phosphorylates the RAPTOR component of the TORC1 complex and inhibits TOR activity (Nukarinen et al., 2016). In plants, N is important for growth but our knowledge about the connections between TOR and N metabolism were hitherto scarce. However, several recent reports have now described the regulations of TORC1 by amino acids in photosynthetic organisms. O’Leary et al. showed that isoleucine or glutamine activate TOR in mature leaves of Arabidopsis (O’Leary et al., 2020). Moreover, loss of function of isopropylmalate synthase 1 (IPMS1), an enzyme involved in Leu synthesis, resulted in the accumulation of Val and to a reduced sensitivity towards TOR inhibitors like AZD-8055 as a consequence of TOR activation (Schaufelberger et al., 2019). In another study, mutation of IPMS1 was also found to be responsible for the Arabidopsis *eva1* mutant phenotype affected in vacuole morphogenesis through hyperactivation of TOR (Cao et al., 2019). As in the previous study, this mutation resulted in the accumulation of Val and other branched-chain amino acids (BCAA) and in TOR activation. Finally, it was found that inorganic N and amino acids activate TOR through the small GTPase Rho-related protein from plants 2 (ROP2) which was also shown to be implicated in auxin induction of TOR activity (Schepetilnikov et al., 2017; Liu et al., 2021). Therefore, the stimulation of TOR activity by amino acids, especially glutamine and BCAA, seems to be a conserved hallmark of this kinase in eukaryotic organisms.

To better establish the relation between N metabolism and the TOR kinase in plants, we describe here the consequences of variations in TOR activity on N metabolism in Arabidopsis. We first investigated the influence of N supply on TOR activity as well as the impact of TOR inhibition on inorganic N uptake and amino acid synthesis. Finally, by using GS inhibitors, we showed that the accumulation of Gln and other amino acids observed after TOR inhibition is dependent on GS activity. Accordingly, growth of TORC1 mutants was improved by GS inhibitors like phosphinothricin. Our results thus provide new evidence of a tight link between the TOR signaling pathway and the N metabolism in plants and show that GS is involved in the accumulation of amino acids observed after TOR inhibition.

## RESULTS

### C/N balance regulates TOR activity

It is now established that TOR is strongly activated by sugar in plants (Xiong et al., 2013; Dobrenel et al., 2016a) and recent studies highlighted the regulation of TOR activity by different nitrogen sources and amino acids such as glutamine (Cao et al., 2019; O’Leary et al., 2020; Liu et al., 2021). Variations in the nutritional environment will affect C/N balance. Therefore, it is important for plants to sense this balance in order to efficiently adapt their metabolism and growth. We thus tried to determine whether the C/N balance could modulate TOR activity in Arabidopsis. In plants, as in other Eukaryotes, TOR phosphorylates the S6 Kinase (S6K) which then phosphorylates the Ribosomal Protein S6 (RPS6) (Mahfouz et al., 2006; Wullschleger et al., 2006). RPS6 phosphorylation on Ser240 has thus been used as a robust readout for TOR activity (Dobrenel et al., 2016b). Using specific antibodies against RPS6 and P-RPS6 we were able to estimate the level of TOR activation by the C/N balance (Figure 1). In the first part of the experiment, 5-days old Arabidopsis seedlings grown on solid medium were transferred for 24 h into liquid medium without sucrose to trigger sugar starvation. The seedlings were then treated for 5h with 1% sucrose in order to stimulate TOR activity +/− 1μM AZD-8055, a potent and validated TOR inhibitor in plants (Montané and Menand, 2013). As expected, the addition of sucrose induces approximatively a 4-fold increase of the P-RPS6/RPS6 ratio, reflecting TOR activation, whereas this activation was abolished when AZD-8055 was added to the medium, despite the presence of sucrose. These expected results validate the experimental setting used to measure the influence of N on TOR activity. In a second experiment, 5-days old Arabidopsis seedlings grown on solid medium were transferred for 24 h into liquid medium without sucrose and KNO_3_. Seedlings were then treated for 5h with or without 1% sucrose and/or 5mM KNO3. The addition of sucrose alone or KNO_3_ alone did not induce TOR activity, but when both nutrients were added to the medium, a 4-fold induction of TOR activity was observed (Figure 1). We also tested 5mM glutamine or ammonium succinate as N sources in combination with sucrose to activate TOR (Supplemental Figure S1). In the same way, TOR activity was higher when both sucrose and a N source like Gln or ammonium were provided. We can thus conclude that both N and sucrose are required for a full activation of TOR.

**Figure 1:**
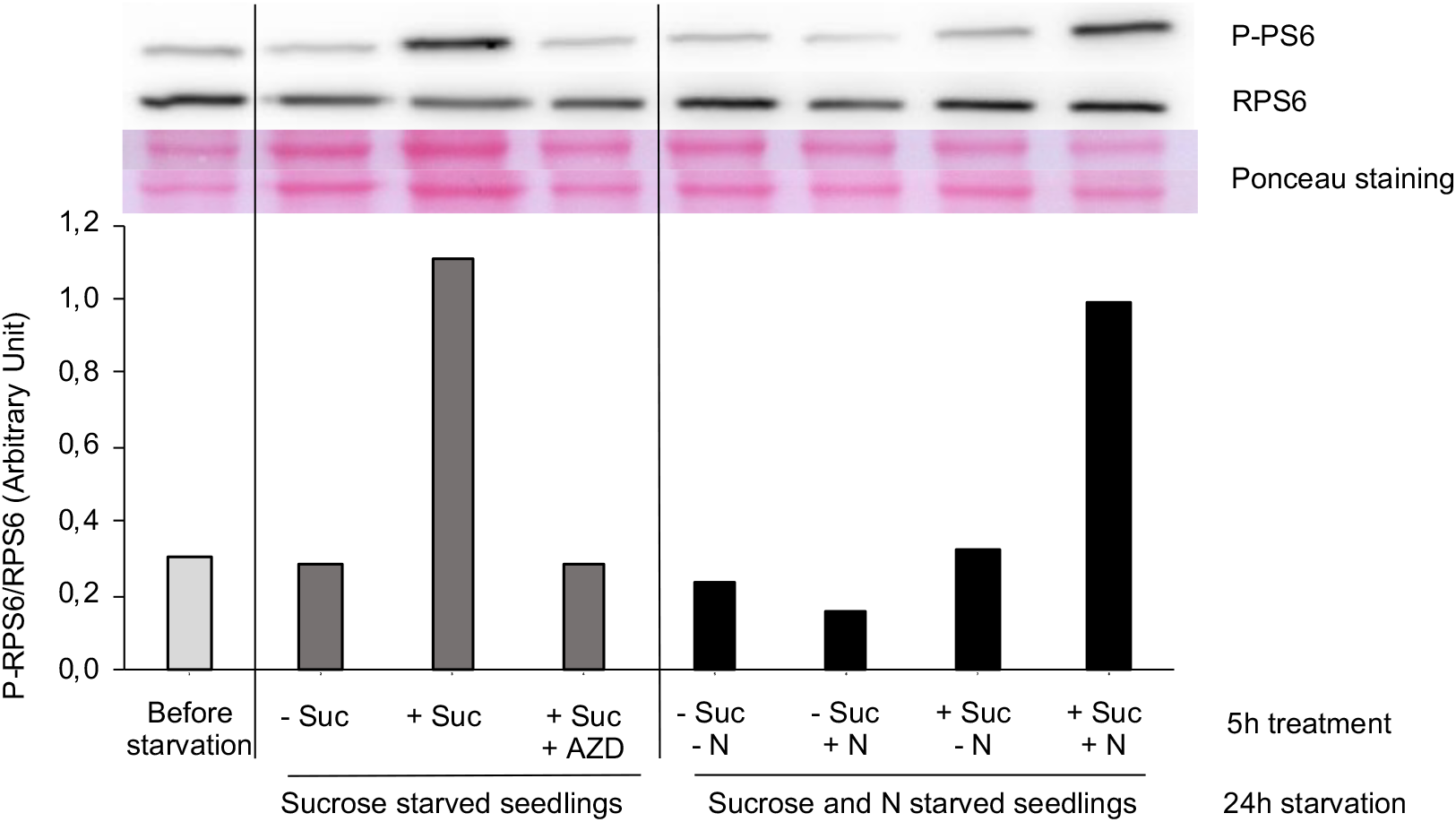
Nitrate is required for the full activation of TOR by sucrose. Five days old seedlings were grown on solid Arabidopsis medium (1mM KNO3 and 0.3% sucrose) then transferred for 24h starvation in liquid Arabidopsis media either without sucrose only, or without sucrose and nitrate. For sucrose starved seedlings (-Suc), 1% Sucrose (+Suc) was then added for 5h with (+AZD) or without 1μM AZD-8055. For sucrose and nitrogen starved seedlings, 1% sucrose (+/− Suc) and/or 5mM KNO3 (+/-N) were added. The same KCl concentration was maintained in the different conditions tested. Twenty five μg of total proteins extracted from around 20 seedlings were loaded for each sample and were used for western blot analysis. Membrane were incubated with phospho-specific antibodies against ribosomal protein S6 (P-RPS6) Ser240 or against total RPS6. Western blots were revealed using Goat antirabbit IgG-HRP as secondary antibody. The Western blot was repeated three times and a typical experiment is shown. A repetition is shown in Supplemental Figure S1. Exposition time: 90sec for P-RPS6, 15 sec for RPS6.

### TORC1 mutants are more sensitive to glutamine

We tried to further define the relations between Gln and TOR by first characterizing the effect of this amino acid on the root growth of TORC1 mutants. To address this question, we used T-DNA insertion mutants affected either in the *Raptor3g* gene (At3g08850, named *raptor 78*) or in the *Lst8-1* gene (At3g18140, named *lst8*). These TORC1 mutants have already been extensively validated and studied (Anderson et al., 2005; Deprost et al., 2005; Moreau et al., 2012; Salem et al., 2018). Mutant plants were grown together with the WT control *in vitro* on increasing Gln concentrations, and primary root growth was measured (Figure 2). The root growth of WT seedlings was slightly but significantly increased by low concentration of Gln compared to the starvation condition, whereas for higher glutamine concentrations (7 to 10 mM), root growth was partially inhibited. The root length of both *raptor 78* and *lst8* mutants was always shorter than WT seedlings and in both mutants root growth was more sensitive to Gln inhibition (Figure 2B). The growth phenotype of *lst8* mutants is generally more severe than the one of *raptor* mutants (Moreau et al., 2012). Coherently, *lst8* mutants seem to be more sensitive to the added Gln with a significative effect on root growth even for low concentrations of Gln when compared to WT and *raptor* mutants. Shoot growth was also more affected by Gln for *raptor 78* and especially *lst8* mutants (Supplemental Figure S2). This higher sensitivity to Gln could be explained by the fact that both *raptor* and *lst8* mutants already contain high levels of this amino acid.

**Figure 2:**
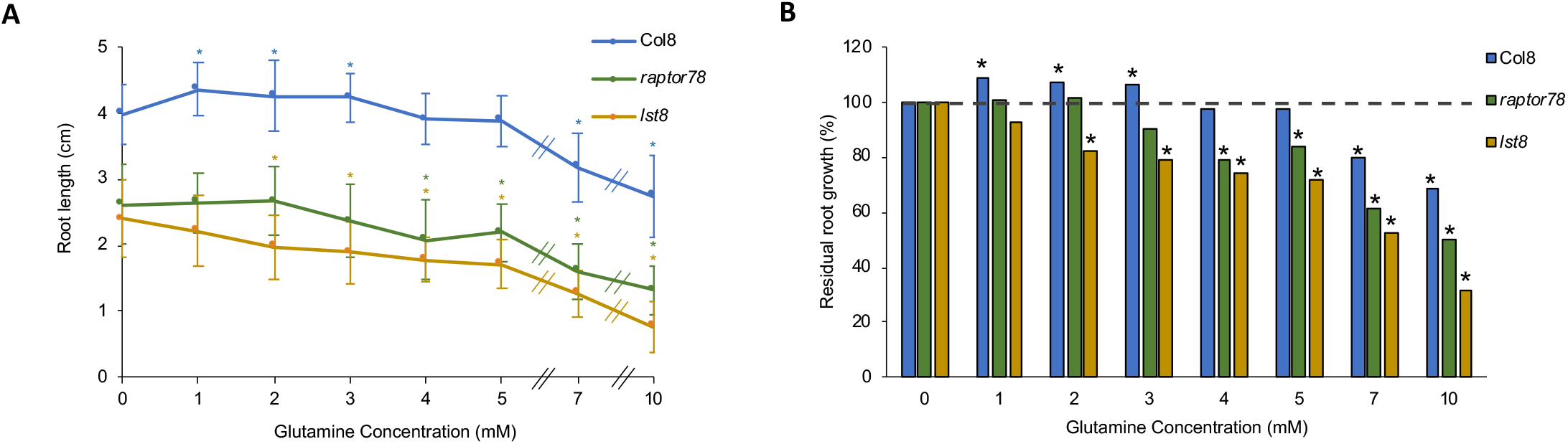
*raptor78* and *lst8* mutants of the TORC1 complex are hypersensitive to glutamine. **(A)** Primary root length and **(B)** Residual root growth of 10 days old wild type control (Col8), raptor78 and lst8 mutants on different glutamine concentrations. Nitrate (0.1mM KNO3) was added for all Gln concentrations to stimulate seed germination. The residual root growth was calculated as a percentage compared to the 0mM glutamine condition (=100%). Results of one typical experiment are shown. n≥12. Stars indicate for each genotype significant differences compared to the control 0mM glutamine condition (Mann and Whitney statistical test, p<0.05).

### TOR inhibition decreases ammonium uptake by the roots

Free Gln and other amino acids can be produced by proteolysis, by ammonium recycling or by the assimilation of inorganic N after uptake from the soil. To elucidate the origin of the amino acid accumulation, and particularly of Gln, when TOR is inhibited, we examined the first step of N assimilation, namely the uptake of nitrate and ammonium by the root system. Depending on the availability of these N sources in the soil, different transporter families are involved in uptake. NPF transporters (Nitrate Transporter 1/ Peptide transporter family) are involved in nitrate uptake for high concentrations (LATS, Low Affinity Transporter System) and NRT2 transporters (Nitrate Transporter 2) are operating at low concentrations (HATS, High Affinity Transporter System) (for review, Kiba and Krapp, 2016). The ammonium transporters’ (AMT/MEP/Rh) superfamily is involved in high affinity ammonium uptake and redistribution within the plant (HATS, for review, von Wirén et al., 2000; Ludewig et al., 2007; Hao et al., 2020). Plant roots also have a low-affinity transport system (LATS) for ammonium. HATS plays a major role when ammonium in the soil is at μM concentration while LATS mainly operates at mM concentrations (Loqué and von Wirén, 2004).

In Chlamydomonas, it has been shown that TOR is involved in the regulation of ammonium uptake, which was found to be higher when cells were treated with rapamycin (Mubeen et al., 2018). To determine if TOR is also involved in the regulation of inorganic N transport in Arabidopsis, we measured ^15^N labelled nitrate or ammonium uptake after 2h of TOR inhibition by AZD-8055 (Supplemental Figure S3A). Ten days old seedlings grown on solid medium were transferred to liquid Arabidopsis media with or without 2μM of AZD-8055. Nitrate uptake was measured with two different ^15^N concentrations, 0.2mM (for HATS) and 5mM (for LATS). TOR inhibition by AZD-8055 had no effect on nitrate uptake for both concentrations (Supplemental Figure S3B), as observed in Chlamydomonas. However, the LATS component of ammonium uptake significantly decreased after TOR inhibition by AZD-8055 whereas the HATS component remained unaffected (Figure 3A). We also measured the LATS component of ammonium uptake after long term TOR inhibition by a lower dose of AZD-8055 and found that it was reduced in a similar way (Supplemental Figure S3C). To confirm this result, we determined ammonium uptake at high concentration (LATS) in *lst8* and *raptor* TORC1 mutants (Figure 3B). As expected, the *raptor* mutants showed a 3-times lower ammonium uptake than WT seedlings, but surprisingly ammonium uptake in *lst8* mutants was at the same level than in WT seedlings. These results suggest that in Arabidopsis TOR is not involved in the regulation of high or low affinity nitrate uptake, but positively regulates ammonium low affinity uptake. Contrary to what was observed in Chlamydomonas by Mubeen et al. (2018), TOR activity seems to stimulate ammonium uptake at high concentrations in a RAPTOR-dependent manner but independently of LST8. This is one of the few examples of a biological process in which RAPTOR and LST8 seem to have different roles.

**Figure 3:**
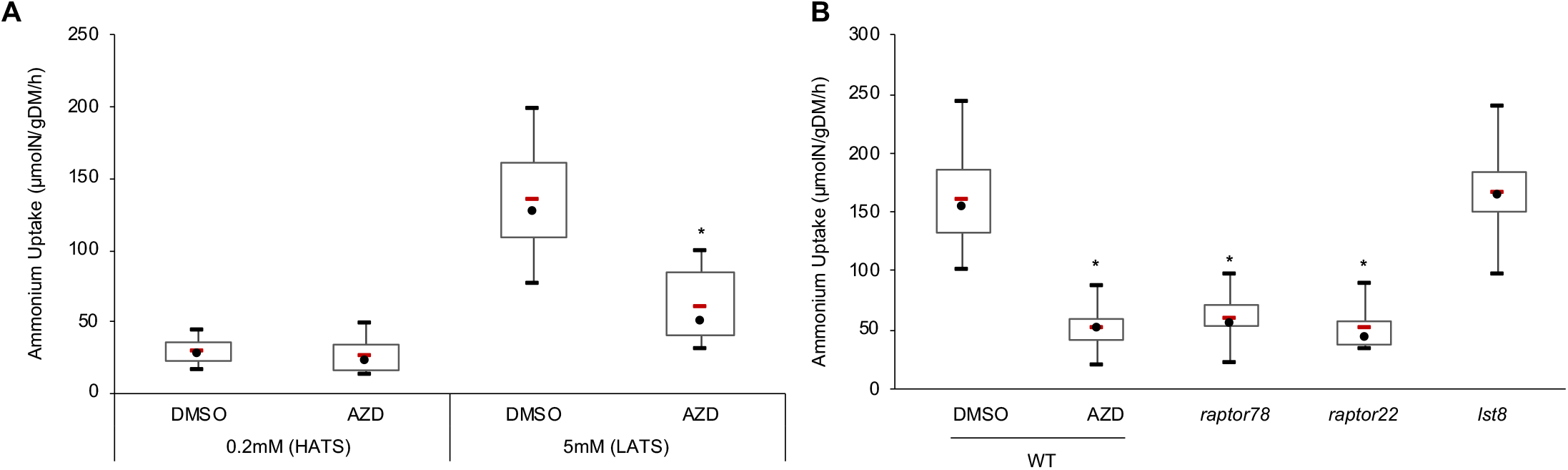
TOR inhibition represses low affinity ammonium uptake in Arabidopsis. Ten days old seedlings were grown on solid Arabidopsis medium (5mM KNO3, 1% sucrose) and treated as described in Supplemental Figure S3 (see also Material and Methods for details). **(A)** Col8 seedlings were pre-treated for 2h with 2μM AZD-8055 before the addition of either 0.2mM ^15^NH_4_ (HATS) or 5mM ^15^NH_4_ (LATS). Plant were collected after 5min and dried before measurements of ^15^N enrichment. n= 18 samples per condition. **(B)** TORC1 mutants (lst8-1, raptor22, raptor78), and WT seedlings were treated with 2μM AZD-8055 in the presence of 5mM ^15^NH_4_ (LATS). n= 14 to 18 (6 samples for raptor22). Red bars represent the mean and black dots represent the median. DMSO are the control vehicle conditions. Stars indicate statistically significant differences compared to WT DMSO condition (non-parametric Kruskal and Wallis Statistical test, p<0.05). Three independent experiments were pooled. gDM: gram dry matter.

### TOR inhibition leads to an ammonium and Gln increase in a GS-dependent manner

The decrease of ammonium uptake observed after TOR inhibition should trigger a lower ammonium and Gln accumulation, whereas it has been extensively documented that TOR inhibition results in Gln accumulation. These seemingly conflicting results prompted us to address the role of GS, the enzyme responsible for Gln synthesis, in the accumulation and growth-inhibitory properties of this amino acid in relation to TOR activity. The GS enzyme catalyzes the amination of glutamate (Glu) using ammonium to produce Gln. The active ingredient of non-selective herbicides like Basta is Glufosinate-Ammonium (GLA), also named phosphinothricin, which is a potent GS inhibitor (Logusch et al., 1990). GLA is a L-glutamate analogue that binds the active site of GS. Methionine-sulfoximine (MSX) is another GS inhibitor that also acts by blocking the GS active site. This GS inhibition, in addition to prevent glutamine production, leads to ammonium accumulation part of which comes from photorespiration and protein degradation (reviewed in Li et al., 2014). To investigate the mechanisms of Gln accumulation after TOR inhibition, we first measured ammonium contents in WT or TORC1 mutant seedlings (Figure 4A). We also used *lst8* mutant suppressed by the introduction of a *yak1* kinase mutation (*yak_lst8*, Forzani et al., 2019). For WT seedlings, GLA addition in the growth medium resulted in a significant increase in ammonium contents compared to DMSO control conditions. This suggests that GLA is active as expected in our experimental conditions. Interestingly, while TOR inhibition represses ammonium uptake, we still observe an increase in ammonium contents after genetic or pharmacological TOR inactivation (Figure 4A). The *raptor* mutation triggers an increase in ammonium concentration which is not augmented by GLA. Conversely, the *lst8* mutants displayed the highest concentration of ammonium while GLA decreased this accumulation. We have already shown before that inactivating the YAK1 kinase suppresses most of the metabolic perturbations brought about by *lst8* mutation (Forzani et al., 2019). However, while ammonium concentration decreased by 50% in *yak_lst8* seedlings, they still contain more ammonium than the WT control.

**Figure 4:**
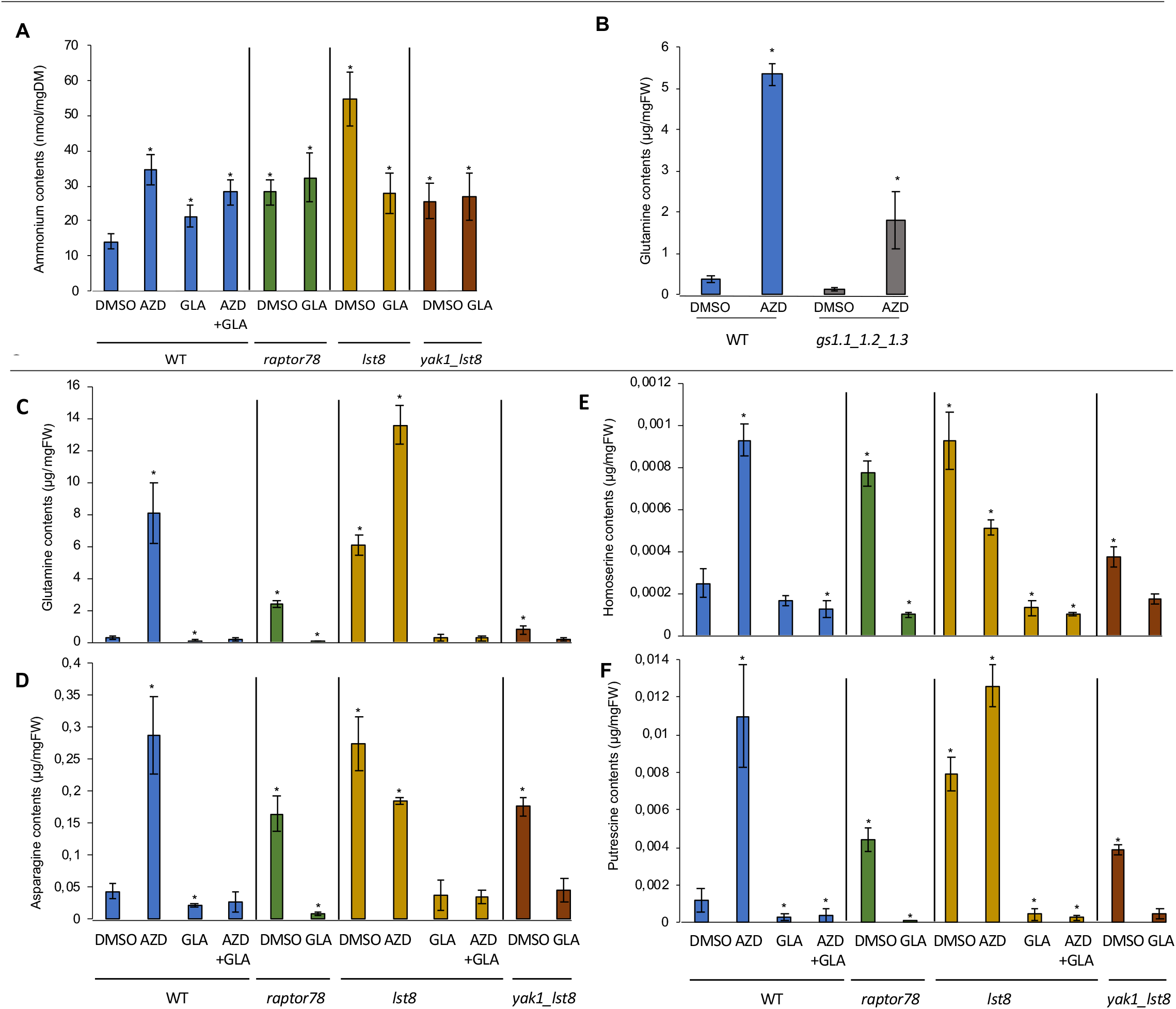
TOR inhibition leads to metabolic changes and Gln accumulation in a GS-dependent manner. Seedlings were grown 5 days on solid Arabidopsis medium then transferred on solid Arabidopsis media with either 0.25μM AZD-8055, or Glufosinate ammonium (GLA) at 2mg/L (GS inhibitor), or both AZD and GLA. The seedlings were harvested after 7 days for metabolomic analysis (n=3). For the experiment with the GS1 triple mutant (B), seedlings were grown directly on 0.25μM AZD-8055 for 10 days. **(A)** Ammonium contents in WT seedlings and TORC1 mutants (raptor78, lst8 and yak1_lst8). Means are the result of 5 independent experiments. **(B)** Effect of AZD-8055 treatment on Gln contents in WT and a GS1 triple mutant (gs1.1×1.2×1.3) (n= 4 and 5 repetitions, for respectively, WT and gs1.1×1.2×1.3 seedlings). **(C, D)** Effect of AZD-8055 or GLA treatments on metabolites contents in WT seedlings and TORC1 mutants. Stars indicate statistically significant differences compared to the WT control plants treated with DMSO condition (non-parametric Mann and Whitney Statistical test, p<0.05).

We then studied the effect of GS inhibition by GLA on the TORC1 mutants. For this purpose, we performed metabolomic analyses using seedlings transferred on medium containing TOR and/or GS inhibitors. Our results confirmed the global accumulation of free amino acids triggered by TOR inhibition that has been observed in earlier reports (Figure 4, B-E; see Ingargiola et al., 2020 and references therein). It is interesting to note that *lst8* mutants treated with AZD-8055 present the strongest accumulation of free Gln compared to control condition (0.25±0.1 μg Gln mg^-1^ fresh weight for WT compared to 13.61±1.2 μg mg^-1^ for *lst8* seedlings on AZD-8055). Without AZD-8055, TORC1 mutants contained more Gln than WT and *lst8* seedlings accumulated two times more Gln than *raptor* mutants (Figure 4C). This higher Gln content in *lst8* mutants could explain their higher sensitivity to the supply of Gln (Figure 2). The pattern of Asn accumulation mirrored that of Gln but at a much lower concentration (Figure 4D), except for the effect of AZD-8055 on *lst8* mutants which lowered Asn concentration but increased Gln. Similarly, TOR deficiency strongly increased the amounts of homoserine (HoSer) (Figure 4E). HoSer is derived from Asp and is an important non proteogenic intermediate amino acid for the production of Met, Thr and Ile. Glu is used for the synthesis of amino acids like GABA, Pro and Arg. Contrary to Gln, TOR inhibition did not significantly affect the level of Glu which often shows a rather constant level in plants. As for the other amino acids, TOR inhibition increased the levels of Arg but also the amounts of both agmatine and putrescine, a polyamine (Figure 4F and Supplemental Figure S4). In Arabidopsis, putrescine is synthesized from Arg via agmatine that is produced by Arg decarboxylase (Hanfrey et al., 2001). Putrescine and other polyamines are involved in the regulation of stress responses and growth (Takahashi 2020). These results show that TOR has a global and strong impact on Gln and Glu metabolisms.

The inhibition of GS activity by GLA treatment strikingly abolished the Gln accumulation caused by TOR inhibition, either after treatment with AZD-8055 or in *raptor* and *lst8* mutants. The levels of other amino acids and polyamines, like Asn, HoSer, agmatine and putrescine were also reduced in TOR-inhibited plants after GLA treatment (Figure 4 and Supplemental Figure S4). As expected, GLA affected Gln contents in control WT plants. In Arabidopsis, the GS enzymes are localized either in the cytosol for the five GS1 isoforms and in the plastids for GS2. To confirm our hypothesis, we used the GS1 triple mutant *gs1.1_1.2_1.3* (Moison et al., 2018). The *gs1.1_1.2_1.3* triple mutant also accumulated Gln after TOR inhibition, but to a lesser extent than the AZD-treated WT plants (Figure 4B). The decrease in Gln level after TOR inhibition was thus lower in the triple *gs1* mutant line than after GLA treatment of WT plants but the GS2 isoenzyme is still active in these mutants compared to GLA-treated plants. Taken together, these data strongly suggest that the Gln accumulation triggered by TOR inhibition is depending on GS activity. Interestingly, GLA also abolished the glucose accumulation resulting from TOR inhibition by AZD-8055 (Supplemental Figure S4). This could indicate a redirection of C fluxes toward organic acid production when GS is inhibited to favor ammonium assimilation.

The observation that GLA suppressed Gln accumulation after TOR inhibition prompted us to determine the effect of GLA on the growth of TORC1 mutants. Five days old TORC1 mutant seedlings were transferred on medium containing different GLA concentrations and the root length was measured 9 days post-transfer (Figure 5). As expected for an herbicide, WT seedlings presented a significant inhibition of shoot and root growth for 2 and 5mg/L of GLA (Supplemental Figure S5A). Indeed, at the highest GLA concentration the WT root length was two-times smaller. However, GLA improved the root growth of TORC1 mutants at low concentrations, especially for *lst8* seedlings and globally TORC1 mutants were less sensitive to GLA than WT (Figure 5 and Supplemental Figure S5A). Similar experiments were performed with MSX, another GS inhibitor, confirming the partial resistance of TORC1 mutants to GS inhibition compared to WT seedlings (Supplemental Figure S5B). The improved root growth of TORC1 mutants after GS inhibition could be the result of the lower level of Gln, or Gln derivatives like putrescine, which are toxic at high concentrations. Surprisingly, the addition of GLA reduced the induction of TOR activity by sucrose (Supplemental Figure S6). This could suggest that the synthesis of Gln is needed to fully induce TOR.

**Figure 5:**
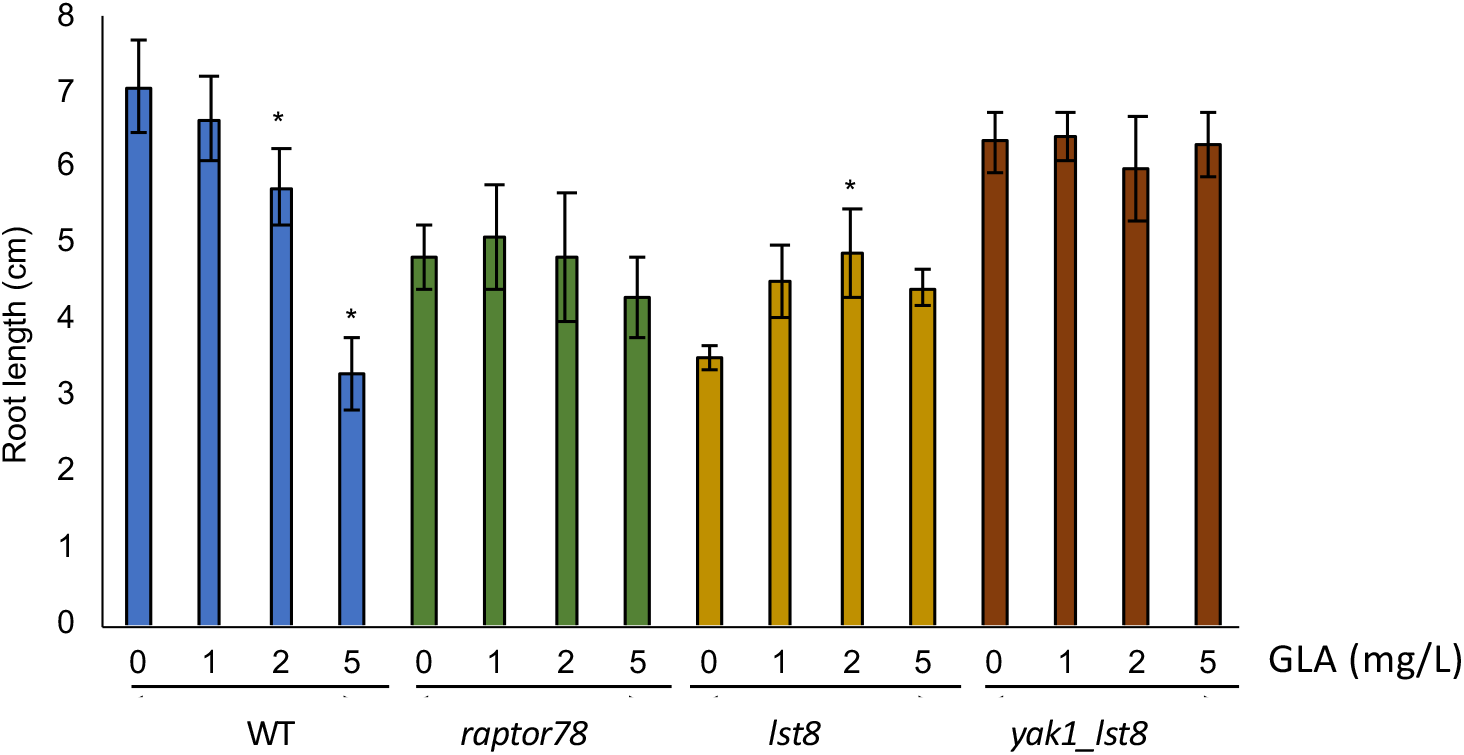
TORC1 mutants are more resistant to GS inhibition by Glufosinate ammonium (GLA) than WT. The root length of WT seedlings and TORC1 mutants (raptor78, lst8, yak_lst8) were measured 9 days posttransfer on Arabidopsis media containing different GLA concentrations. This figure shows one representative experiment among 4 experiments. Stars indicate significant differences compared to the control 0 mg/L condition for each genotype (n=20, non-parametric Kruskal and Wallis Statistical test, p<0.05).

Conversely the *gs1* triple mutant, as expected since they have less GS activity and consequently contain less Gln, were hypersensitive to GLA compared to WT (Supplemental Figure S7). Coherently, this *gs1* mutant, as well as the quadruple *amt* (ammonium transporter) mutant, are more resistant to TOR inhibition than WT when grown on AZD-8055 (Supplemental Figure S8). This result suggests that a reduction in Gln content correlates with a decrease in the sensitivity to TOR inhibition.

### TOR inhibition leads to a deregulation of N metabolism and GS enzymes in Arabidopsis

To gain more insight into the regulations of Gln metabolism by TOR in Arabidopsis, we asked how the inhibition of TOR is affecting the mRNA level of genes involved in ammonium uptake and nitrate assimilation. We thus analyzed by quantitative RT-PCR the expression of some marker genes either in the presence of nitrate or ammonium (Figure 6A). On nitrate, several genes were more expressed after TOR inhibition such as *AMT1;1* (ammonium transporter), *GLN1;1, GLN1;3* (encoding GS1 isoforms) or *GLU2* (a Fd-GOGAT (ferredoxin-dependent Gln 2 oxo-glutamate amino transferase) mainly expressed in roots (Coschigano et al., 1998)). More genes were induced by TOR inhibition when plants were grown on ammonium, including *AMT1;3, AMT1;5, Nii* (nitrite reductase), *Nia1* and *Nia2* (nitrate reductase) as well as *GLN1;1-1;3* and *GLU2*. The stimulation in gene expression was usually higher on ammonium and even more in the *lst8* mutant. To focus on Gln metabolism, we also determined GS protein levels and activities after TOR inhibition (Figure 6, B-C). Nitrate-grown seedlings treated with AZD had a lower GS activity compared to the WT (Figure 6C). Accordingly, GS activity was also lower in *raptor78* and *lst8* mutants whereas the *yak_lst8* suppressor mutant displayed a GS activity equivalent to the WT. The fact that pharmacological and genetical TORC1 inhibitions decrease GS activity is somewhat counter-intuitive given the massive Gln accumulation resulting from these inhibitions. Finally, we analysed GS proteins level by immuno-blot in 10 days old seedlings (Figure 6B and Supplemental Figure S9). The *gs1.1_1.2_1.3* mutant presents an important decrease of GS1 protein level compared to the WT, accompanied by an increase of GS2 protein. This compensatory mechanism was already described before for this mutant line (Moison et al., 2018). Genetic or pharmacologic TOR inhibition leads to an increase of GS1 and GS2 protein levels. The *raptor* and *lst8* mutants display a strong increase of GS protein levels which is higher than in seedings treated by AZD-8055. These results suggest that the GS protein is more abundant, but less active after TOR inhibition and thus that TOR may be required for GS activation.

**Figure 6:**
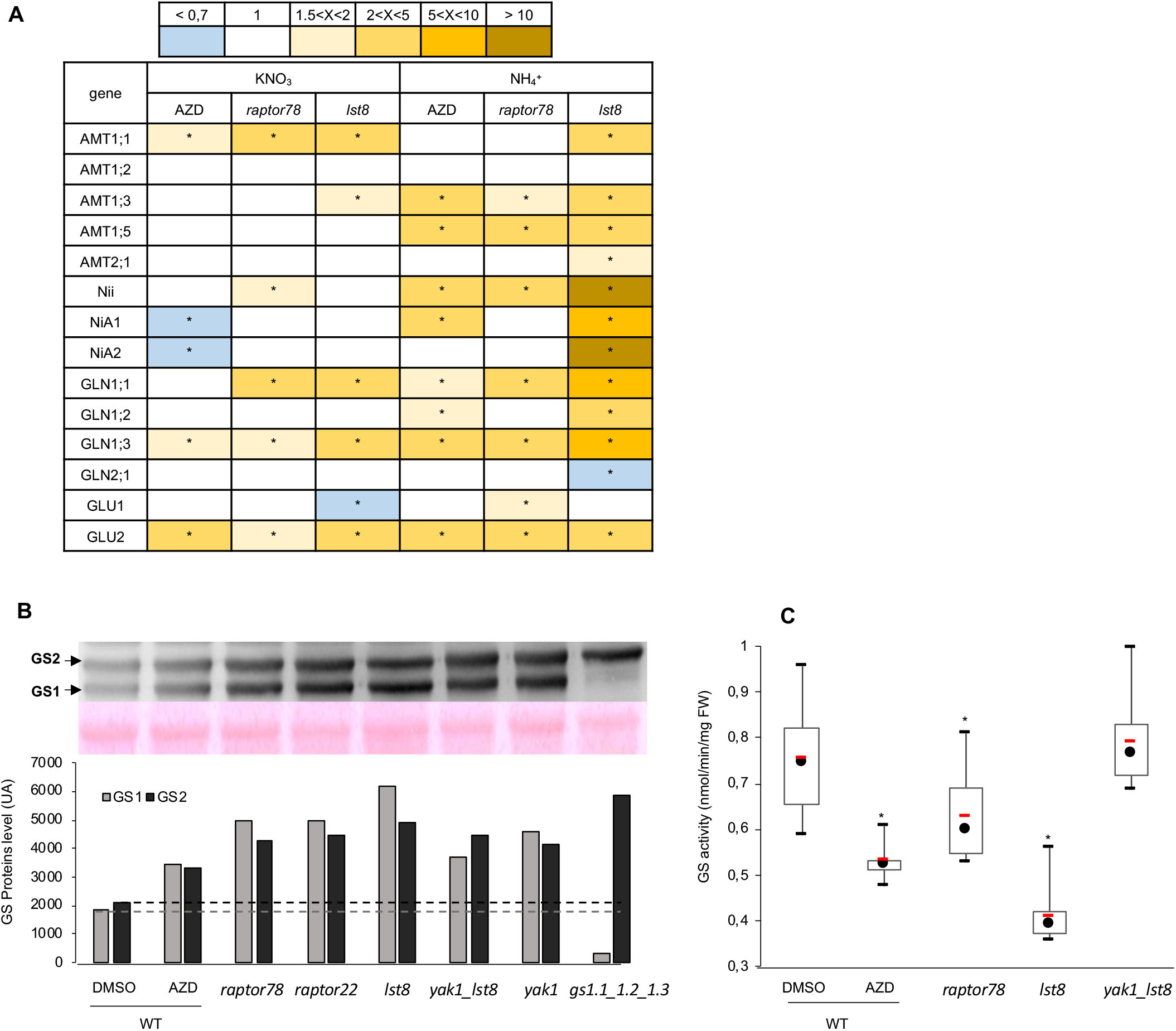
Impact of TOR inhibition on the expression of N metabolism genes and GS activity. (A) Effects of genetic or pharmacological TOR inhibition on the expression of N assimilation genes, AMT: ammonium transporters, Nii: nitrite reductase, Nia: nitrate reductase; GLN: Gln synthetase, GLU: Glu synthase. Gene expression levels were determined by quantitative RT-PCR using 10 days old WT and TORC1 mutants seedlings (raptor78 and lst8) grown on 5mM KNO3 or ammonium succinate. The changes in gene expression were determined as described in Material and Methods with expression levels in nitrate or ammonium-grown plants treated with either AZD-8055 (0.25μM) or DMSO as controls. Mean ratios from three independent experiments are shown as fold changes. Stars indicate statistical difference (Mann & Whitney statistical test). (B) GS1 and GS2 amounts were determined using WT seedlings, TORC1 mutants (raptor22, raptor78, lst8, yak1 and yak1_lst8) and the gs1.1_1.2_1.3 triple mutant grown on Arabidopsis medium with 5mM KNO3 and 1% sucrose. Western Blot was performed with a total GS antibody (1:1000), 25 μg of total proteins were loaded for each sample extracted from 19 to 35 seedlings. GS1: 38-40 kDa, GS2: 47.4 kDa. Western blot signals were quantified by densitometry. (C) GS activity in 10 days old WT seedlings and TORC1 mutants (raptor78, lst8 and yak1_lst8) grown as above (the results are the means of 6 independent experiments with 19 to 35 seedlings). Red bars represent the mean and black dots represent the median. Stars indicate significant differences compared to WT (non-parametric Mann & Whitney Statistical test, p<0.05).

## DISCUSSION

The intricate connections between the availability of nutrients and the regulations of energy-demanding cellular processes such as mRNA translation or cell growth must be tightly and precisely orchestrated. A wealth of studies in animals and yeast have shown that the conserved TOR kinase lies at the heart of these interactions (Efeyan et al., 2015; González and Hall, 2017). Several studies have also demonstrated that in algae and plants TOR activity is strongly influenced by the N status and the supply of amino acids (Mubeen et al., 2018; Cao et al., 2019; O’Leary et al., 2020; Upadhyaya et al., 2020; Liu et al., 2021). The main goal of this study was to better describe and understand the large accumulation of free amino acids frequently observed in both plants and algae after TOR inhibition (see Ingargiola et al., 2020 and references therein).

The first objective was to determine the effect of different nutrients and N sources on TOR activity in Arabidopsis. To estimate TOR activity, we used as a proxy the phosphorylation of RPS6 on Ser240 (Dobrenel et al., 2016b). We confirm that, as previously observed, sucrose is a very potent inducer of TOR activity in Arabidopsis (Xiong et al., 2013; Dobrenel et al., 2016b). However, both N and C supplies seem to be required for TOR activation (Figure 1). This result is in line with the previous observations that both sugar and nitrate are necessary to coordinate the synthesis of the translational machinery, which is largely TOR-dependent (Dobrenel et al., 2016b; Scarpin et al. 2020), and that mRNA translation is highly sensitive to the C/N status of the plant (Guttiérez et al., 2007). Ammonium and Gln were also found to activate TOR in sugar starved Arabidopsis seedlings re-supplied with sucrose (Supplemental Figure S1). The induction of TOR activity by Gln was already recently described in several reports (for example O’Leary et al., 2020; Liu et al., 2021). Coherently, the inhibition of Gln synthesis by the herbicide glufosinate (GLA), that blocks GS activity, suppresses the stimulation of TOR activity by sucrose (Supplemental Figure S6). These data suggest that Gln, originating from GS activity, has a dual role with respect to TOR activity: synthesis of Gln seems to be needed for TOR activation by sucrose while its accumulation appears correlated with sensitivity to TOR inhibition.

We focused our study on the accumulation of Gln, which is by far the most affected amino acid after TOR inhibition (Moreau et al., 2012; Ren et al., 2012; Caldana et al., 2013; Salem et al., 2018; Forzani et al., 2019). Indeed, Gln concentration increased by more than 50 times in *lst8* mutants (Figure 4). However, this concerns only free Gln and not the total amount of proteins in the plants, level of which decreases upon TOR inhibition (Deprost et al., 2007). This Gln accumulation could be due to (at least) three mechanisms which are (i) an increase in glutamine biosynthesis from ammonium by GS, (ii) a decrease in the use of Gln for protein or amino acid synthesis by transaminases like GOGAT and/or (iii) an increase in protein degradation and recycling phenomena. For the latter explanation, it is indeed known that TOR inhibition triggers autophagic processes leading to protein degradation within the vacuole (Liu and Bassham, 2010). Similarly, TOR inhibition decreases mRNA translation and thus the use of free amino acids (Deprost et al., 2007). Nevertheless, these processes are slow and Gln accumulates very rapidly after TOR inhibition (Mubeen et al., 2018). Moreover, although the quantities of most amino acids augment after TOR inhibition, Gln levels show by far the highest increase while one would expect a similar accumulation of all amino acids, reflecting the overall composition of proteins, if mRNA translation is stopped. Most amino acids derive, directly or indirectly, from Gln by transamination reactions and would probably not accumulate if the use of Gln is diminished. This probably rules out the second hypothesis for Gln accumulation.

Indeed, the report by Mubeen et al. (2018) showing that ^15^N incorporation into amino acids was higher in cells treated by rapamycin supports the assumption that a higher N assimilation is the cause for the build-up of Gln. The authors also found that ammonium uptake was higher after TOR inhibition by rapamycin. The fact that Gln is also the amino acid which is used for transport between organs in Arabidopsis could also explain the fact that it is accumulated to higher extend than other amino acids (Dinkeloo et al., 2018). To further explore this hypothesis of deregulation of Gln synthesis, we first determined the impact of TOR inhibition on nitrate and ammonium transport using ^15^N labelling. Surprisingly, nitrate uptake was not significantly affected by TOR inhibition, although its LATS component was reduced by AZD-8055 treatment (Supplemental Figure S3). Conversely, the low affinity/high-capacity transport system of ammonium ions was decreased after either pharmacological or genetic inhibition of TOR (Figure 3). AMT transporters are involved in the HATS component of ammonium transport (Liu and von Wirén, 2017) and it has been suggested earlier that the LATS ammonium transport might occur via the potassium channel (Wang et al., 1994). Interestingly, it was shown recently that inhibition of TOR reduces potassium uptake in Arabidopsis (Deng et al., 2020). In yeast, TOR regulates potassium homeostasis through the control of the plasma membrane (H+)-ATPase (Mahmoud et al., 2017). This decrease in low affinity ammonium uptake was also observed after long term TOR inhibition (Supplemental Figure S3). At high concentrations, ammonium in its uncharged NH_3_ form can diffuse through the membrane with the help of aquaporins. Moreover, a role for aquaporins of the TIP family (tonoplast intrinsic proteins) in ammonium accumulation was also described (Loqué et al., 2005). It is therefore difficult to identify the TOR-dependent targets which mediate the decrease in ammonium uptake after TOR inhibition.

Conversely, ammonium concentration increased in the same conditions either after pharmacological or genetic inhibition of TOR (Figure 4A). A similar build-up of ammonium was detected in Chlamydomonas treated with rapamycin (Mubeen et al., 2018). In yeast, TOR inhibition results in the up-regulation of the NPR1 kinase activity and in the subsequent phosphorylation and activation of the MEP2 ammonium transporter (Boeckstaens et al., 2014). But yeast uses ammonium as a poor N source and activates its transport and assimilation in N starved conditions. Interestingly, the tonoplast-localized receptor-like kinase CAP1, mutation of which results in higher ammonium concentrations in the cytosol and lower uptake through the tonoplast, complemented the yeast *npr1* mutants, thus suggesting that TOR may control ammonium homeostasis in plants through CAP1 (Bai et al., 2014). The fact that TOR-inhibited plants accumulate ammonium and Gln whereas uptake is reduced could suggest that ammonium originates from another source than from the growth medium. It could for example be envisaged that TOR inhibition stimulates the photorespiratory pathway thereby increasing ammonium production by Gly oxidation in mitochondria. Indeed, a large proportion of ammonium is coming from photorespiration in C3 plants like Arabidopsis.

A recent screen for ammonium-insensitive Arabidopsis mutants identified the GS2 gene as a causal mutation and the authors suggested that higher GS2 activity in high ammonium conditions produced more H+ and resulted in an acid stress response inhibiting growth (Hachiya et al., 2021). This could be also the case after TOR inhibition in Arabidopsis since Gln is accumulating in a GS-dependent manner (Figure 4). Furthermore, in yeast rapamycin treatment has been shown to inhibit the plasma membrane (H+)-ATPase (Mahmoud et al., 2017). If this inhibition of H+ efflux also happens in plants, it would further contribute to cytosol acidification and growth reduction. Coherently, we discovered that inhibition of total GS activity by the herbicide GLA drastically reduced Gln and other amino acids accumulation (Figure 4). This result shows that Gln is originating from ammonium assimilation by GS. Since GLA inhibits both cytosolic GS1 and chloroplastic GS2, we cannot conclude on the specific contribution of GS isoforms on Gln accumulation (Takano and Dayan, 2020). GLA is very specific for GS and competes with Glu for the active site. GLA is phosphorylated by GS into phospho-L-phosphinothricin, but ammonium cannot be later incorporated to produce Gln. This leads to an irreversible inactivation of GS and to a strong reduction in the overall enzyme activity since GLA is readily translocated within the plant (Takano and Dayan, 2020). Therefore, the reduction in Gln accumulation is probably the cause for the growth improvement observed in TORC1 mutants treated with GLA, whereas growth of WT plants was hampered by this herbicide as expected (Figure 5). However, *lst8* mutants and AZD-treated WT seedlings displayed a marked reduction in ammonium accumulation after treatment with GLA whereas ammonium levels increased in treated control plants (Figure 4). Indeed, the action of GLA as herbicide is thought to be through ammonium accumulation resulting from the impairment of GS activity. However, other cellular processes such as photorespiration could be affected and it has been shown that GLA increases the cellular pool of Leu and Val which are known inducers of TOR activity in plants (Wendler et al., 1990; Cao et al., 2019).

We also showed that TORC1 mutants were hypersensitive to high concentrations of Gln in the growth medium compared to WT seedlings, growth of which was also affected for concentrations above 5 mM (Figure 2). In consequence, it is probably both ammonium buildup and assimilation in addition to Gln accumulation which are toxic when TOR activity is reduced. However, at this point it is difficult to conclude that Gln *per se*, or a molecule derived from Gln, is the cause of the growth reduction in TOR-limited plants. For example, Arg is synthesized from Gln and Glu and is catabolized into agmatine which then serves as a precursor for the synthesis of putrescine, a precursor for longer polyamines like spermidine and spermine. In Arabidopsis it seems that most putrescine is synthesized from Arg unlike other plants where ornithine is the precursor molecule (Hanfrey et al., 2001). While Arg levels are only modestly increased by TOR inhibition, the levels of both agmatine and putrescine were highly augmented (Supplemental Figure S4). High levels of putrescine have been shown to inhibit growth but putrescine and agmatine amounted to 100 times less than Gln in TOR-inhibited plants (Fuell et al., 2010). Very recently, it was suggested that TOR activity regulates polyamine metabolism and degradation by controlling the translation of the involved enzymes (Salazar-Diaz et al., 2021). Similarly, Salem et al. (2018) have also measured increased levels of Gln, agmatine, and putrescine in Arabidopsis *raptor* mutants.

We have also observed in ammonium-fed *lst8* mutants or in plants treated with AZD-8055 an increase in the mRNA level of enzymes involved in nitrate assimilation as well as of the AMT ammonium transporters (Figure 6). It was shown earlier that excess Gln repress the expression of the *Amt1;1* transporter (Rawat et al., 1999). Conversely, we observed an increase in *Amt1;1* expression in ammonium-grown *lst8* mutants which accumulate Gln. On the contrary, the *Nii* and *Nia* genes showed the expected repression by Gln in *lst8* mutants (Vincentz et al., 1993). While GS1/GS2 mRNA and protein levels generally increased, total GS activity was found to decrease following pharmacological or genetic inhibition of TOR (Figure 6). This suggests that the higher Gln accumulation could trigger, by a feedback mechanism, a lower GS activity, as well as lower ammonium uptake and concomitantly an increase in GS expression to compensate for this decrease of enzymatic activity. The fact that the decrease in GS activity seems to be proportional to Gln levels (Figure 6) would support this hypothesis. Alternatively, GS could be activated by TOR through a direct or indirect phosphorylation. Indeed, it is known that GS can be regulated by phosphorylation (Ji et al., 2019). Accordingly, the GOGAT enzyme was found to be phosphorylated in a TOR-dependent manner (van Leene et al., 2019). However, rapamycin treatment of Chlamydomonas cells resulted in higher GS activity (Mubeen et al., 2018). These differences may reflect the different ways of life between an unicellular alga for which excess Gln can be easily excreted into the surrounding water and a multicellularland plant.

## CONCLUSION

Our data reveal that the Gln and amino acid accumulation caused by TOR inhibition is suppressed by genetic or pharmacological inhibition of GS activity. Furthermore, GS inhibitor such as the herbicide GLA clearly improves the growth of TORC1 mutants. The study of N fluxes within the plant with heavy isotope tracer will be required to obtain a more complete view of the metabolic perturbations responsible for the effect of TOR and/or GS inhibition on metabolism and growth of plants.

## MATERIALS AND METHODS

### Plant Material

In this study, we used *Arabidopsis thaliana* ecotype Columbia 8 as wild-type. The *lst8* mutant (noted *lst8-1*, At3g18140, SALK_002459) was obtained from the Nottingham Arabidopsis Stock Center (NASC). The *raptor* mutants (At3g08850) noted *raptor78* (SALK_078159) and *raptor22* (SALK_022096) were obtained from the NASC or the Arabidopsis Biological Resource Center (ABRC). The *yak1* (SALK_131306) and *yak1_lst8* double mutant were previously described (Forzani et al., 2019). The triple *gs1.1_1.2_1.3* mutant was kindly provided by Dr. Céline Masclaux-Daubresse. This mutant was previously described in Moison et al., 2018. The quadruple *amt* mutant line (*amt1;1 amt1;2 amt1;3 amt2;1*) was described in Yuan et al. (2007).

### Plant Growth Conditions

Seedlings were grown *in vitro* on vertical plates containing Arabidopsis medium with 5mM KNO_3_, 1% sucrose, 2.5mM KH_2_PO_4_, 2mM MgSO_4_, 1mMCaCl_2_, 0.05% 2-(N-Morpholino) ethanesulfonic acid sodium salt (MES), 1X microelements (Moreau et al., 2012), 5mM Iron Ammonium Citrate and 0.5% Phytagel. The seeds were stratified at 4°C for 2 days and then transferred in controlled growth chambers at 21°C with a photoperiod of 16h light/8h night. For phenotypic analysis, root length was measured with the NeuronJ plugin in the ImageJ software.

### Protein extraction, Western Blot Analysis and TOR activity measurements

For TOR activity measurements, 5 days old seedlings were grown on solid Arabidopsis medium (1mM KNO_3_ and 0,3% sucrose) and then transferred for 24h starvation in liquid Arabidopsis media either only without sucrose, or without sucrose and nitrate. For sucrose starved seedlings, medium supplemented with 1% Sucrose (+Suc) was then added for 5h with or without 1μM AZD-8055 (Sellekchem). For sucrose and nitrogen starved seedlings, media supplemented with +/− 1% sucrose (+/− Suc) +/− 5mM KNO_3_ (+/-N) were added. The level of RPS6 phosphorylation was used as a readout of TOR activity. Total proteins were extracted from whole seedlings using 1X Laemmli Buffer (50mM Tris-HCl pH6.8, 2% SDS, 10% Glycerol, 0.0025% bromophenol blue) and 400mM DTT. Protein extracts were heated 5 min at 95°C and then centrifuged to removed cell debris. Protein concentrations were determined using Bradford reagent (Bio-Rad) at 595nm. An equal amount of proteins (25μg for TOR activity measurement, 28μg for GS protein levels) were loaded in 12% acrylamide resolving SDS-PAGE gel and separated by migration for 1h at 150V and 2A. Proteins were then transferred on PVDF membranes following the Trans-Blot®Turbo™Blotting System protocol. Membranes were blocked with 5% milk-PBS solution (13.7mM NaCl, 0.27mM KCl, 1mM Na_2_HPO_4_, 0.2mM KH_2_PO_4_) and 0,05% Tween20. Then, membranes were incubated with rabbit primary antibodies directed against either total RPS6, or P-RPS6 (for both dilution 1:5000) as described in Dobrenel et al. (2016b), or with anti-GS antibodies (dilution 1/1000) (Lothier et al., 2011). Goat anti-rabbit IgG-HRP (Sigma) was used as secondary antibody (dilution 1/5000). Immunodetection was performed using the Clarity Western ECL detection kit (Bio-Rad). The equal transfer of proteins on the membrane was evaluated by Ponceau Staining. P-RPS6/RPS6 ratio and GS protein levels were quantified with the ImageJ software.

### ^15^N Uptake Analysis

Ten days old seedlings were pre-treated in 1mM KNO_3_ +/− AZD 2μM for 2h (Supplemental Figure S2A) and were after washed 1 min in 0.1mM CaSO_4_ solution, treated 5 min in liquid Arabidopsis media containing as N source 0.2mM (for HATS) or 5mM (for LATS) K^15^NO_3_ or (^15^NH_4_)2SO_4_ (99% labelling). Afterwards the seedlings were washed again 1 min in the CaSO_4_ solution. After 48h in oven (70°C), ^15^N uptake was determined by isotope ratio mass spectrometry from 0.6 to 1mg of dry matter. The uptake was then calculated as:

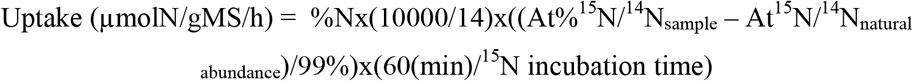

### Metabolomic Analysis

For metabolomic analysis and ammonium measurements, 5 days old seedlings were transferred for 7 days on new media with DMSO as a control, 0,25μM AZD-8055, 2mg/L glufosinate ammonium (Sigma) or 0,25μM AZD-8055 + 2mg/L glufosinate ammonium. Glutamine and metabolites levels were determined by GC-MS on *in vitro* grown whole seedlings. For details on sample preparations, metabolite quantifications and analyses see Forzani et al. (2019).

### Ammonium measurement

For ammonium measurements, 500μL of 80% ethanol was added to 100mg of fresh material then incubated 1hr at 4°C, and centrifuged at 15000 rpm (4°C for 15 min). Supernatants were collected and conserved for the analysis. Two 80% ethanol extraction were further performed (500μL then 250μL), followed by a 50% ethanol extraction and then water extraction (250μL). 2mM NH_4_Cl solution was used for building a reference curve with a range between 0 and 200nmol. Ammonium was then measured as described by Moison et al. (2018).

### RNA extraction and quantitative RT-PCR

Total RNAs were extracted from 10 days whole seedlings (around 50mg of fresh material) using the Trizol^®^ reagent method. Total RNA (1μg) was used for DNAse I Thermos Scientific treatment and RT reaction using RevertAid H Minus RT Thermo Scientific. The quantitative RT-PCR was performed using pink Sybr from Biorad. The expression of the Actin2 gene (At3g18780) and EF1 gene (At5g60390) were used as genes reference. The primers used for these analyses are listed in Supplemental Table S1.

### Glutamine Synthetase activity measurement

GS activity was evaluated by the measurement of γ-glutamylhydroxamate at 540nm. Enzymes were extracted from whole seedlings (150mg of fresh material) with an extraction buffer containing 10mM Na-EDTA, 10mM MgCl2 and 250mM pH7,6 Tris-HCl. β-mercaptoethanol (13 μM final concentration) and 4 nM leupeptine were added in this buffer before extraction. Samples were centrifuged at 12 000 rpm for 10 min at 4°C and the supernatants were collected. GS activity was then determined as described previously (Moreau et al., 2012).

## Acknowledgements

We wish to thank Dr. Céline Masclaux-Daubresse for providing seeds of the *gs1* triple mutant (*gs1.1×1.2×1.3*) and GS antibodies, Dr. Nicolaus von Wirén for the quadruple *qko* AMT seedlings, Virginie Bréhaut for *Nii* and *Nia* primers and Anne Krapp, Thomas Girin, Sylvie Ferrario-Méry, Camila Caldana, Guillaume Pilot, and Christophe Robaglia for inspiring discussions on N metabolism and the TOR kinase. We also thank our colleagues at IJPB for discussions and advices.

## Supplemental Data

**FigS1: Nitrate is required for the full activation of TOR by sucrose and other sources of N, such as Gln or ammonium, also allow full activation of TOR by sucrose.**

**FigS2: Effect of increasing Gln concentration on the growth of TORC1 mutants.**

**FigS3: ^15^N influx experiments.**

**FigS4: Metabolite levels are affected by TOR inhibition.**

**FigS5: Effect of increasing concentration of the herbicides glufosinate (GLA) and methionine sulfoximine (MSX) on the growth of TORC1 mutants.**

**FigS6: Treatment with the GS inhibitor glufosinate (GLA) decreases TOR induction by sucrose.**

**FigS7: Root growth of the triple *gs1* mutant is hypersensitive to Glufosinate ammonium (GLA) compared to the WT.**

**FigS8: Root growth of GS and ammonium transporters (AMT) mutants is less sensitive to TOR inhibition.**

**FigS9: Additional replicate of Western blot analysis for GS1 and GS2 proteins (Figure 6).**

**Supplemental Table S1: List of the primers used in Figure 6.**

## Parsed Citations

**Adami A. García-Alvarez B, Arias-Palomo E, Barford D, Llorca O (2007) Structure of TOR and its complex with KOG1. Mol Cell 27: 509-516**

Google Scholar: <u>Author Only</u> <u>Title Only</u> <u>Author and Title</u>

**Anderson G, Veit B, Hanson M (2005) The Arabidopsis AtRaptor genes are essential for post-embryonic plant growth. BMC Biol 3: 12**

Google Scholar: <u>Author Only</u> <u>Title Only</u> <u>Author and Title</u>

**Aylett CH, Sauer E, Imseng S, Boehringer D, Hall MN, Ban N, Maier T (2016) Architecture of human mTOR complex 1. Science 351: 48-52**

Google Scholar: <u>Author Only</u> <u>Title Only</u> <u>Author and Title</u>

**Bai L, Ma X, Zhang G, Song S, Zhou Y, Gao L, Miao Y, Song CP (2014) A Receptor-Like Kinase Mediates Ammonium Homeostasis and Is Important for the Polar Growth of Root Hairs in Arabidopsis. Plant Cell 26: 1497-1511**

Google Scholar: <u>Author Only</u> <u>Title Only</u> <u>Author and Title</u>

**Barrada A. Djendli M, Desnos T, Mercier R, Robaglia C, Montané MH, Menand B (2019) A TOR-YAK1 signaling axis controls cell cycle, meristem activity and plant growth in Arabidopsis. Development 146: 3**

Google Scholar: <u>Author Only</u> <u>Title Only</u> <u>Author and Title</u>

**Beck T, Hall MN (1999) The TOR signalling pathway controls nuclear localization of nutrient-regulated transcription factors. Nature 402: 689-692**

Google Scholar: <u>Author Only</u> <u>Title Only</u> <u>Author and Title</u>

**Bloom AJ, Sukrapanna SS, Warner RL (1992) Root respiration associated with ammonium and nitrate absorption and assimilation by barley. Plant Physiol 99: 1294-1301**

Google Scholar: <u>Author Only</u> <u>Title Only</u> <u>Author and Title</u>

**Boeckstaens M, Llinares E, Van Vooren P, Marini AM (2014) The TORC1 effector kinase Npr1 fine tunes the inherent activity of the Mep2 ammonium transport protein. Nat Commun 5: 3101**

Google Scholar: <u>Author Only</u> <u>Title Only</u> <u>Author and Title</u>

**Brunkard JO (2020) Exaptive Evolution of Target of Rapamycin Signaling in Multicellular Eukaryotes. Dev Cell 54: 142-155**

Google Scholar: <u>Author Only</u> <u>Title Only</u> <u>Author and Title</u>

**Caldana C, Li Y, Leisse A. Zhang Y, Bartholomaeus L, Fernie AR, Willmitzer L, Giavalisco P (2013) Systemic analysis of inducible target of rapamycin mutants reveal a general metabolic switch controlling growth in Arabidopsis thaliana. Plant J 73: 897-909**

Google Scholar: <u>Author Only</u> <u>Title Only</u> <u>Author and Title</u>

**Cao P, Kim SJ, Xing A. Schenck CA, Liu L, Jiang N, Wang J, Last RL, Brandizzi F (2019) Homeostasis of branched-chain amino acids is critical for the activity of TOR signaling in. Elife 8**

Google Scholar: <u>Author Only</u> <u>Title Only</u> <u>Author and Title</u>

**Coschigano KT, Melo-Oliveira R, Lim J, Coruzzi GM (1998) Arabidopsis gls mutants and distinct Fd-GOGAT genes. Implications for photorespiration and primary nitrogen assimilation. Plant Cell 10: 741-752**

Google Scholar: <u>Author Only</u> <u>Title Only</u> <u>Author and Title</u>

**Crespo JL, Powers T, Fowler B, Hall MN (2002) The TOR-controlled transcription activators GLN3, RTG1, and RTG3 are regulated in response to intracellular levels of glutamine. Proc Natl Acad Sci U S A 99: 6784-6789**

Google Scholar: <u>Author Only</u> <u>Title Only</u> <u>Author and Title</u>

**Deng K, Wang W, Feng L, Yin H, Xiong F, Ren M (2020) Target of rapamycin regulates potassium uptake in Arabidopsis and potato. Plant Physiol Biochem 155: 357-366**

Google Scholar: <u>Author Only</u> <u>Title Only</u> <u>Author and Title</u>

**Deprost D, Truong H, Robaglia C, Meyer C (2005) An Arabidopsis homolog of RAPTOR/KOG1 is essential for early embryo development. Biochem Biophys Res Commun 326: 844-850**

Google Scholar: <u>Author Only</u> <u>Title Only</u> <u>Author and Title</u>

**Deprost D, Yao L, Sormani R, Moreau M, Leterreux G, Nicolaï M, Bedu M, Robaglia C, Meyer C (2007) The Arabidopsis TOR kinase links plant growth, yield, stress resistance and mRNA translation. EMBO Rep 8: 864-870**

Google Scholar: <u>Author Only</u> <u>Title Only</u> <u>Author and Title</u>

**Dinkeloo K, Boyd S, Pilot G (2018) Update on amino acid transporter functions and on possible amino acid sensing mechanisms in plants. Semin Cell Dev Biol 74: 105-113**

Google Scholar: <u>Author Only</u> <u>Title Only</u> <u>Author and Title</u>

**Dobrenel T, Caldana C, Hanson J, Robaglia C, Vincentz M, Veit B, Meyer C (2016) TOR Signaling and Nutrient Sensing. Annu Rev**

**Plant Biol 67: 261-285**

Google Scholar: <u>Author Only</u> <u>Title Only</u> <u>Author and Title</u>

**Dobrenel T, Mancera-Martínez E, Forzani C, Azzopardi M, Davanture M, Moreau M, Schepetilnikov M, Chicher J, Langella O, Zivy M, Robaglia C, Ryabova LA, Hanson J, Meyer C (2016) The Arabidopsis TOR Kinase Specifically Regulates the Expression of Nuclear Genes Coding for Plastidic Ribosomal Proteins and the Phosphorylation of the Cytosolic Ribosomal Protein S6. Front Plant Sci 7: 1611**

Google Scholar: <u>Author Only</u> <u>Title Only</u> <u>Author and Title</u>

**Díaz-Troya S, Pérez-Pérez ME, Pérez-Martín M, Moes S, Jeno P, Florencio FJ, Crespo JL (2011) Inhibition of protein synthesis by TOR inactivation revealed a conserved regulatory mechanism of the BiP chaperone in Chlamydomonas. Plant Physiol 157: 730-741**

Google Scholar: <u>Author Only</u> <u>Title Only</u> <u>Author and Title</u>

**Efeyan A. Comb WC, Sabatini DM (2015) Nutrient-sensing mechanisms and pathways. Nature 517: 302-310**

Google Scholar: <u>Author Only</u> <u>Title Only</u> <u>Author and Title</u>

**Fichtner F, Dissanayake IM, Lacombe B, Barbier F (2021) Sugar and Nitrate Sensing: A Multi-Billion-Year Story. Trends Plant Sci 26: 352-374**

Google Scholar: <u>Author Only</u> <u>Title Only</u> <u>Author and Title</u>

**Forzani C, Duarte GT, Van Leene J, Clément G, Huguet S, Paysant-Le-Roux C, Mercier R, De Jaeger G, Leprince AS, Meyer C (2019) Mutations of the AtYAK1 Kinase Suppress TOR Deficiency in Arabidopsis. Cell Rep 27: 3696-3708**

Google Scholar: <u>Author Only</u> <u>Title Only</u> <u>Author and Title</u>

**Fuell C, Elliott KA, Hanfrey CC, Franceschetti M, Michael AJ (2010) Polyamine biosynthetic diversity in plants and algae. Plant Physiol Biochem 48: 513-520**

Google Scholar: <u>Author Only</u> <u>Title Only</u> <u>Author and Title</u>

**González A, Hall MN (2017) Nutrient sensing and TOR signaling in yeast and mammals. EMBO J 36: 397-408**

Google Scholar: <u>Author Only</u> <u>Title Only</u> <u>Author and Title</u>

**Guan M, de Bang TC, Pedersen C, Schjoerring JK (2016) Cytosolic Glutamine Synthetase Gln1;2 Is the Main Isozyme Contributing to GS1 Activity and Can Be Up-Regulated to Relieve Ammonium Toxicity. Plant Physiol 171: 1921-1933**

Google Scholar: <u>Author Only</u> <u>Title Only</u> <u>Author and Title</u>

**Gutiérrez RA, Lejay LV, Dean A, Chiaromonte F, Shasha DE, Coruzzi GM (2007) Qualitative network models and genome-wide expression data define carbon/nitrogen-responsive molecular machines in Arabidopsis. Genome Biol 8: R7**

Google Scholar: <u>Author Only</u> <u>Title Only</u> <u>Author and Title</u>

**Hachiya T, Inaba J, Wakazaki M, Sato M, Toyooka K, Miyagi A, Kawai-Yamada M, Sugiura D, Nakagawa T, Kiba T, Gojon A, Sakakibara H (2021) Excessive ammonium assimilation by plastidic glutamine synthetase causes ammonium toxicity in Arabidopsis thaliana. Nat Commun 12: 4944**

Google Scholar: <u>Author Only</u> <u>Title Only</u> <u>Author and Title</u>

**Hanfrey C, Sommer S, Mayer MJ, Burtin D, Michael AJ (2001) Arabidopsis polyamine biosynthesis: absence of ornithine decarboxylase and the mechanism of arginine decarboxylase activity. Plant J 27: 551-560**

Google Scholar: <u>Author Only</u> <u>Title Only</u> <u>Author and Title</u>

**Hao DL, Zhou JY, Yang SY, Qi W, Yang KJ, Su YH (2020) Function and Regulation of Ammonium Transporters in Plants. Int J Mol Sci 21**

Google Scholar: <u>Author Only</u> <u>Title Only</u> <u>Author and Title</u>

**Ingargiola C, Turqueto Duarte G, Robaglia C, Leprince AS, Meyer C (2020) The Plant Target of Rapamycin: A Conduc TOR of Nutrition and Metabolism in Photosynthetic Organisms. Genes (Basel) 11: 1285**

Google Scholar: <u>Author Only</u> <u>Title Only</u> <u>Author and Title</u>

**Ji Y, Li Q, Liu G, Selvaraj G, Zheng Z, Zou J, Wei Y (2019) Roles of Cytosolic Glutamine Synthetases in Arabidopsis Development and Stress Responses. Plant Cell Physiol 60: 657-671**

Google Scholar: <u>Author Only</u> <u>Title Only</u> <u>Author and Title</u>

**Kiba T, Krapp A (2016) Plant Nitrogen Acquisition Under Low Availability: Regulation of Uptake and Root Architecture. Plant Cell Physiol 57: 707-714**

Google Scholar: <u>Author Only</u> <u>Title Only</u> <u>Author and Title</u>

**Krapp A (2015) Plant nitrogen assimilation and its regulation: a complex puzzle with missing pieces. Curr Opin Plant Biol 25: 115-122**

Google Scholar: <u>Author Only</u> <u>Title Only</u> <u>Author and Title</u>

**Lee P, Cho BR, Joo HS, Hahn JS (2008) Yeast Yak1 kinase, a bridge between PKA and stress-responsive transcription factors,**

**Hsf1 and Msn2/Msn4. Mol Microbiol 70: 882-895**

Google Scholar: <u>Author Only</u> <u>Title Only</u> <u>Author and Title</u>

**Li B, Li G, Kronzucker HJ, Baluška F, Shi W (2014) Ammonium stress in Arabidopsis: signaling, genetic loci, and physiological targets. Trends Plant Sci 19: 107-114**

Google Scholar: <u>Author Only</u> <u>Title Only</u> <u>Author and Title</u>

**Liu GY, Sabatini DM (2020) mTOR at the nexus of nutrition, growth, ageing and disease. Nat Rev Mol Cell Biol 21: 183-203**

Google Scholar: <u>Author Only</u> <u>Title Only</u> <u>Author and Title</u>

**Liu Y, Bassham D (2010) TOR is a negative regulator of autophagy in Arabidopsis thaliana. PLoS One 5: e11883**

Google Scholar: <u>Author Only</u> <u>Title Only</u> <u>Author and Title</u>

**Liu Y, Duan X, Zhao X, Ding W, Wang Y, Xiong Y (2021) Diverse nitrogen signals activate convergent ROP2-TOR signaling in Arabidopsis. Dev Cell 56: 1283-1295**

Google Scholar: <u>Author Only</u> <u>Title Only</u> <u>Author and Title</u>

**Liu Y, von Wirén N (2017) Ammonium as a signal for physiological and morphological responses in plants. J Exp Bot 68: 2581-2592** Google Scholar: <u>Author Only</u> <u>Title Only</u> <u>Author and Title</u>

**Logusch EW, Walker DM, McDonald JF, Franz JE, Villafranca JJ, DiIanni CL, Colanduoni JA, Li B, Schineller JB (1990) Inhibition of Escherichia coli glutamine synthetase by alpha- and gamma-substituted phosphinothricins. Biochemistry 29: 366-372**

Google Scholar: <u>Author Only</u> <u>Title Only</u> <u>Author and Title</u>

**Loqué D, von Wirén N (2004) Regulatory levels for the transport of ammonium in plant roots. J Exp Bot 55:1293-305.**

Google Scholar: <u>Author Only</u> <u>Title Only</u> <u>Author and Title</u>

**Lothier J, Gaufichon L, Sormani R, Lemaître T, Azzopardi M, Morin H, Chardon F, Reisdorf-Cren M, Avice JC, Masclaux-Daubresse C (2011) The cytosolic glutamine synthetase GLN1;2 plays a role in the control of plant growth and ammonium homeostasis in Arabidopsis rosettes when nitrate supply is not limiting. J Exp Bot 62: 1375-1390**

Google Scholar: <u>Author Only</u> <u>Title Only</u> <u>Author and Title</u>

**Ludewig U, Neuhäuser B, Dynowski M (2007) Molecular mechanisms of ammonium transport and accumulation in plants. FEBS Lett 581: 2301-2308**

Google Scholar: <u>Author Only</u> <u>Title Only</u> <u>Author and Title</u>

**Maegawa K, Takii R, Ushimaru T, Kozaki A (2015) Evolutionary conservation of TORC1 components, TOR, Raptor, and LST8, between rice and yeast. Mol Genet Genomics 290: 2019-2030**

Google Scholar: <u>Author Only</u> <u>Title Only</u> <u>Author and Title</u>

**Mahfouz MM, Kim S, Delauney AJ, Verma DP (2006) Arabidopsis TARGET OF RAPAMYCIN interacts with RAPTOR, which regulates the activity of S6 kinase in response to osmotic stress signals. Plant Cell 18: 477-490**

Google Scholar: <u>Author Only</u> <u>Title Only</u> <u>Author and Title</u>

**Mahmoud S, Planes MD, Cabedo M, Trujillo C, Rienzo A, Caballero-Molada M, Sharma SC, Montesinos C, Mulet JM, Serrano R (2017) TOR complex 1 regulates the yeast plasma membrane proton pump and pH and potassium homeostasis. FEBS Lett 591: 1993-2002**

Google Scholar: <u>Author Only</u> <u>Title Only</u> <u>Author and Title</u>

**Martinez-Outschoorn UE, Peiris-Pagés M, Pestell RG, Sotgia F, Lisanti MP (2017) Cancer metabolism: a therapeutic perspective. Nat Rev Clin Oncol 14: 113**

Google Scholar: <u>Author Only</u> <u>Title Only</u> <u>Author and Title</u>

**Menand B, Desnos T, Nussaume L, Berger F, Bouchez D, Meyer C, Robaglia C (2002) Expression and disruption of the Arabidopsis TOR (target of rapamycin) gene. Proc Natl Acad Sci U S A 99: 6422-6427**

Google Scholar: <u>Author Only</u> <u>Title Only</u> <u>Author and Title</u>

**Meng D, Yang Q, Wang H, Melick CH, Navlani R, Frank AR, Jewell JL (2020) Glutamine and asparagine activate mTORC1 independently of Rag GTPases. J Biol Chem 295: 2890-2899**

Google Scholar: <u>Author Only</u> <u>Title Only</u> <u>Author and Title</u>

**Moison M, Marmagne A, Dinant S, Soulay F, Azzopardi M, Lothier J, Citerne S, Morin H, Legay N, Chardon F, Avice JC, Reisdorf-Cren M, Masclaux-Daubresse C (2018) Three cytosolic glutamine synthetase isoforms localized in different-order veins act together for N remobilization and seed filling in Arabidopsis. J Exp Bot 69: 4379-4393**

Google Scholar: <u>Author Only</u> <u>Title Only</u> <u>Author and Title</u>

**Montané MH, Menand B (2013) ATP-competitive mTOR kinase inhibitors delay plant growth by triggering early differentiation of meristematic cells but no developmental patterning change. J Exp Bot 64: 4361-4374**

Google Scholar: <u>Author Only</u> <u>Title Only</u> <u>Author and Title</u>

**Moreau M, Azzopardi M, Clément G, Dobrenel T, Marchive C, Renne C, Martin-Magniette ML, Taconnat L, Renou JP, Robaglia C,**

**Meyer C (2012) Mutations in the Arabidopsis homolog of LST8/GβL, a partner of the target of Rapamycin kinase, impair plant growth, flowering, and metabolic adaptation to long days. Plant Cell 24: 463-481**

Google Scholar: <u>Author Only</u> <u>Title Only</u> <u>Author and Title</u>

**Mubeen U, Jüppner J, Alpers J, Hincha DK, Giavalisco P (2018) Target of Rapamycin Inhibition in. Plant Cell 30: 2240-2254**

Google Scholar: <u>Author Only</u> <u>Title Only</u> <u>Author and Title</u>

**Nukarinen E, Nägele T, Pedrotti L, Wurzinger B, Mair A, Landgraf R, Börnke F, Hanson J, Teige M, Baena-Gonzalez E, Dröge-Laser W, Weckwerth W (2016) Quantitative phosphoproteomics reveals the role of the AMPK plant ortholog SnRK1 as a metabolic master regulator under energy deprivation. Sci. Rep. 6, 31697**

Google Scholar: <u>Author Only</u> <u>Title Only</u> <u>Author and Title</u>

**O’Brien J A, Vega A, Bouguyon E, Krouk G, Gojon A, Coruzzi G, Gutiérrez RA (2016) Nitrate transport, sensing and responses in plants. Molecular Plant 9: 837-856**

Google Scholar: <u>Author Only</u> <u>Title Only</u> <u>Author and Title</u>

**O’Leary BM, Oh GGK, Lee CP, Millar AH (2020) Metabolite Regulatory Interactions Control Plant Respiratory Metabolism via Target of Rapamycin (TOR) Kinase Activation. Plant Cell 32: 666-682**

Google Scholar: <u>Author Only</u> <u>Title Only</u> <u>Author and Title</u>

**Pérez-Pérez ME, Couso I, Crespo JL (2017) The TOR Signaling Network in the Model Unicellular Green Alga Chlamydomonas reinhardtii. Biomolecules 7**

Google Scholar: <u>Author Only</u> <u>Title Only</u> <u>Author and Title</u>

**Rawat SR, Silim SN, Kronzucker HJ, Siddiqi MY, Glass AD (1999) AtAMT1 gene expression and NH4+ uptake in roots of Arabidopsis thaliana: evidence for regulation by root glutamine levels. Plant J 19: 143-152**

Google Scholar: <u>Author Only</u> <u>Title Only</u> <u>Author and Title</u>

**Ren M, Venglat P, Qiu S, Feng L, Cao Y, Wang E, Xiang D, Wang J, Alexander D, Chalivendra S, Logan D, Mattoo A, Selvaraj G, Datla R (2012) Target of rapamycin signaling regulates metabolism, growth, and life span in Arabidopsis. Plant Cell 24: 4850-4874**

Google Scholar: <u>Author Only</u> <u>Title Only</u> <u>Author and Title</u>

**Salazar-Díaz K, Dong Y, Papdi C, Ferruzca-Rubio EM, Olea-Badillo G, Ryabova LA, Dinkova TD (2021) TOR senses and regulates spermidine metabolism during seedling establishment and growth in maize and Arabidopsis. iScience 24: 103260**

Google Scholar: <u>Author Only</u> <u>Title Only</u> <u>Author and Title</u>

**Salem MA, Li Y, Bajdzienko K, Fisahn J, Watanabe M, Hoefgen R, Schöttler MA, Giavalisco P (2018) RAPTOR Controls Developmental Growth Transitions by Altering the Hormonal and Metabolic Balance. Plant Physiol 177: 565-593**

Google Scholar: <u>Author Only</u> <u>Title Only</u> <u>Author and Title</u>

**Scarpin MR, Leiboff S, Brunkard JO (2020) Parallel global profiling of plant TOR dynamics reveals a conserved role for LARP1 in translation. Elife 9**

Google Scholar: <u>Author Only</u> <u>Title Only</u> <u>Author and Title</u>

**Schaufelberger M, Galbier F, Herger A, de Brito Francisco R, Roffler S, Clement G, Diet A, Hörtensteiner S, Wicker T, Ringli C (2019) Mutations in the Arabidopsis ROL17/isopropylmalate synthase 1 locus alter amino acid content, modify the TOR network, and suppress the root hair cell development mutant lrx1. J Exp Bot 70: 2313-2323**

Google Scholar: <u>Author Only</u> <u>Title Only</u> <u>Author and Title</u>

**Schepetilnikov M, Makarian J, Srour O, Geldreich A, Yang Z, Chicher J, Hammann P, Ryabova LA (2017) GTPase ROP2 binds and promotes activation of target of rapamycin, TOR, in response to auxin. EMBO J 36: 886-903**

Google Scholar: <u>Author Only</u> <u>Title Only</u> <u>Author and Title</u>

**Soto-Burgos J, Bassham DC (2017) SnRK1 activates autophagy via the TOR signaling pathway in Arabidopsis thaliana. PLoS One 12: e0182591**

Google Scholar: <u>Author Only</u> <u>Title Only</u> <u>Author and Title</u>

**Stracka D, Jozefczuk S, Rudroff F, Sauer U, Hall MN (2014) Nitrogen source activates TOR (target of rapamycin) complex 1 via glutamine and independently of Gtr/Rag proteins. J Biol Chem 289: 25010-25020**

Google Scholar: <u>Author Only</u> <u>Title Only</u> <u>Author and Title</u>

**Takahashi T (2020) Plant Polyamines. Plants 9: 511**

Google Scholar: <u>Author Only</u> <u>Title Only</u> <u>Author and Title</u>

**Takano HK, Dayan FE (2020) Glufosinate-ammonium: a review of the current state of knowledge. Pest Manag Sci 76: 3911-3925**

Google Scholar: <u>Author Only</u> <u>Title Only</u> <u>Author and Title</u>

**Tanigawa M, Maeda T (2017) An In Vitro TORC1 Kinase Assay That Recapitulates the Gtr-Independent Glutamine-Responsive TORC1 Activation Mechanism on Yeast Vacuoles. Mol Biol Cell 37: 21**

Google Scholar: <u>Author Only</u> <u>Title Only</u> <u>Author and Title</u>

**Tate JJ, Rai R, De Virgilio C, Cooper TG (2021) N- and C-terminal Gln3-Tor1 interaction sites: one acting negatively and the other positively to regulate nuclear Gln3 localization. Genetics 217: 4**

Google Scholar: <u>Author Only</u> <u>Title Only</u> <u>Author and Title</u>

**Upadhyaya S, Agrawal S, Gorakshakar A, Rao BJ (2020) TOR kinase activity in Chlamydomonas reinhardtii is modulated by cellular metabolic states. FEBS Lett 594: 3122-3141**

Google Scholar: <u>Author Only</u> <u>Title Only</u> <u>Author and Title</u>

**Van Leene J, Han C, Gadeyne A, Eeckhout D, Matthijs C, Cannoot B, De Winne N, Persiau G, Van De Slijke E, Van de Cotte B, Stes E, Van Bel M, Storme V, Impens F, Gevaert K, Vandepoele K, De Smet I, De Jaeger G (2019) Capturing the phosphorylation and protein interaction landscape of the plant TOR kinase. Nat Plants 5: 316-327**

Google Scholar: <u>Author Only</u> <u>Title Only</u> <u>Author and Title</u>

**Vidal EA, Alvarez JM, Araus V, Riveras E, Brooks MD, Krouk G, Ruffel S, Lejay L, Crawford NM, Coruzzi GM, Gutiérrez RA (2020) Nitrate in 2020: Thirty Years from Transport to Signaling Networks. Plant Cell 32: 2094-2119**

Google Scholar: <u>Author Only</u> <u>Title Only</u> <u>Author and Title</u>

**Villar VH, Merhi F, Djavaheri-Mergny M, Durán RV (2015) Glutaminolysis and autophagy in cancer. Autophagy 11: 1198-1208**

Google Scholar: <u>Author Only</u> <u>Title Only</u> <u>Author and Title</u>

**von Wirén N, Gazzarrini S, Gojon A, Frommer WB (2000) The molecular physiology of ammonium uptake and retrieval. Curr Opin Plant Biol 3: 254-261**

Google Scholar: <u>Author Only</u> <u>Title Only</u> <u>Author and Title</u>

**Vincentz M, Moureaux T, Leydecker MT, Vaucheret H, Caboche M (1993) Regulation of nitrate and nitrite reductase expression in Nicotiana plumbaginifolia leaves by nitrogen and carbon metabolites. Plant J 3: 315-324**

Google Scholar: <u>Author Only</u> <u>Title Only</u> <u>Author and Title</u>

**Wang MY, Glass A, Shaff JE, Kochian LV (1994) Ammonium Uptake by Rice Roots (III. Electrophysiology). Plant Physiol 104: 899-906**

Google Scholar: <u>Author Only</u> <u>Title Only</u> <u>Author and Title</u>

**Wendler C, Barniske M, Wild A (1990) Effect of phosphinothricin (glufosinate) on photosynthesis and photorespiration of C3 and C 4 plants. Photosynth Res 24: 55-61**

Google Scholar: <u>Author Only</u> <u>Title Only</u> <u>Author and Title</u>

**Wullschleger S, Loewith R, Hall MN (2006) TOR signaling in growth and metabolism. Cell 124: 471-484**

Google Scholar: <u>Author Only</u> <u>Title Only</u> <u>Author and Title</u>

**Xiong Y, McCormack M, Li L, Hall Q, Xiang C, Sheen J (2013) Glucose-TOR signalling reprograms the transcriptome and activates meristems. Nature 496: 181-186**

Google Scholar: <u>Author Only</u> <u>Title Only</u> <u>Author and Title</u>

**Yuan L, Loqué D, Kojima S, Rauch S, Ishiyama K, Inoue E, Takahashi H, von Wirén N (2007) The organization of high-affinity ammonium uptake in Arabidopsis roots depends on the spatial arrangement and biochemical properties of AMT1-type transporters. Plant Cell 19:2636-2652**

Google Scholar: <u>Author Only</u> <u>Title Only</u> <u>Author and Title</u>

**Yuan L, Sheng X, Willson AK, Roque DR, Stine JE, Guo H, Jones HM, Zhou C, Bae-Jump VL (2015) Glutamine promotes ovarian cancer cell proliferation through the mTOR/S6 pathway. Endocr Relat Cancer 22: 577-591**

Google Scholar: <u>Author Only</u> <u>Title Only</u> <u>Author and Title</u>

